# E2F1 promotes, JNK and DIAP1 inhibit, and chromosomal position has little effect on radiation-induced Loss of Heterozygosity in *Drosophila*

**DOI:** 10.1101/2023.05.09.540084

**Authors:** Jeremy Brown, Tin Tin Su

## Abstract

Loss of Heterozygosity (LOH) can occur when a heterozygous mutant cell loses the remaining wild type allele to become a homozygous mutant. LOH can have physiological consequences if, for example, the affected gene encodes a tumor suppressor. We used two fluorescent reporters to study mechanisms of LOH induction by X-rays, a type of ionizing radiation (IR), in *Drosophila* larval wing discs. IR is used to treat more than half of cancer patients, so understanding its effects is of biomedical relevance. Quantitative analysis of IR-induced LOH at seven different positions between the telomere and the centromere, the first such study that we know of, showed that while a homologous chromosome is required to produce LOH after irradiation, position along the chromosome makes little difference. This suggests that IR-induced LOH occurs via non-recombination-based mechanisms, unlike what was previously reported for spontaneously occurring LOH. Using a focused screen, we identified E2F1 as a key promotor of LOH and further testing suggests a mechanism involving its role in cell cycle regulation rather than apoptosis. We leveraged the loss of a transcriptional repressor through LOH to express transgenes specifically in cells that have already acquired LOH. This approach identified JNK signaling and apoptosis as key determinants of LOH maintenance. These studies reveal previously unknown mechanisms for generation and elimination of cells with chromosome aberrations after exposure to IR.

**Summary:** Loss of Heterozygosity (LOH) can lead to homozygous mutant cells, with potential physiological consequences if it affects tumor suppressor genes. Using fluorescent reporters of LOH in *Drosophila* wing discs, we found that X-ray-induced LOH requires a homologous chromosome but was unaffected by position along the chromosome. This suggests non-recombination mechanisms for X-ray-induced LOH, unlike spontaneous LOH. We found that E2F1 promotes LOH while JNK signaling and apoptosis eliminates cells with LOH, uncovering new insights into chromosomal aberrations post-exposure to radiation.

## Introduction

Loss of Heterozygosity (LOH) occurs when a heterozygous cell loses an allele to become homozygous. LOH can have physiological consequences if, for example, the lost allele encodes a tumor suppressor. Possible mechanisms for LOH include mutations in the gene, aneuploidy of whole or segments of chromosomes, and mitotic recombination between the locus and the centromere (producing -/- and +/+ twin spots from a -/+ cell). We are using *Drosophila melanogaster* as a model to study LOH that follows exposure to X-rays, a type of ionizing radiation (IR). IR causes DNA strand breaks and its ability to induce apoptosis is the reason IR is used to treat half of cancer patients. Understanding IR’s effects on genome integrity is a crucial step towards rendering radiation therapy safe.

Classical and recent studies of IR-induced LOH in *Drosophila* employed visible adult markers such as *multiple wing hair* (*mwh*) and *forked (f)* that alter the number or morphology of actin-based hairs on the cell surface (Baker *et al.* 1978). Starting with heterozygotes of recessive alleles, induced homozygosity is detectable on a cell-by-cell basis. In these studies, exposure to IR occurred in the larval stages while the resulting LOH was scored many days later in the adult. To be able to monitor steps in between, we have been developing fluorescent LOH reporters that can be used at all developmental stages.

In our published work (Brown *et al.* 2020), we used the QF/QS module wherein QF binds its recognition sequence QUAS to promote transcription while QS represses QF (Riabinina *et al.* 2015). psc-QF drives the expression of QUAS-tdTomato (Tom) throughout the larval wing disc (Fig 1A) unless repressed by ubiquitously-expressed QS^9B^ inserted at 100E1 on chromosome 3 (Fig 1F). Loss of QS^9B^ by LOH results in de-repression of QF and cell-autonomous Tom expression (Fig 1C-C’). Our published study produced a surprising result; IR-induced *mwh* LOH was lower by more than 10-fold compared to IR-induced QS^9B^ LOH detected under identical experimental conditions in adult wings (Brown *et al.* 2020). Classical studies suggest that segmental aneuploidy (loss of a chromosome segment) is the primary mechanism for IR-induced LOH (Baker *et al.* 1978). If breakage and loss of the corresponding chromosome tip is indeed the mechanism, *mwh* LOH would accompany the loss of more genes compared to QS^9B^ LOH because *mwh* is further from the telomere than QS^9B^ (Fig 1F). Resulting *mwh* LOH cells may be less fit than QS^9B^ LOH cells, explaining the difference in IR-induced LOH incidence we saw.

**Fig 1.**
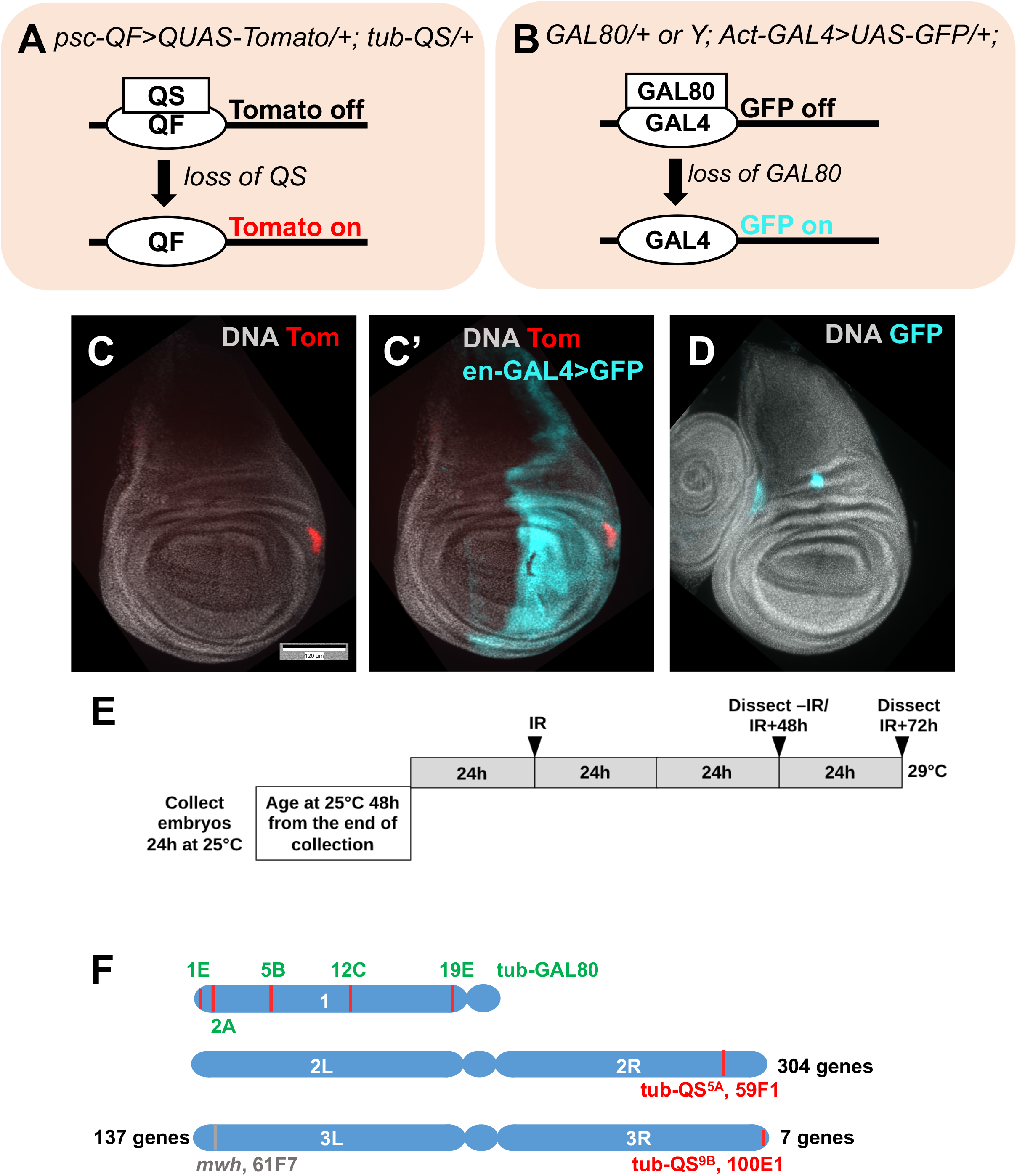
Fluorescent reporters for loss-of-heterozygosity. A. Loss of QS results in de-repression of psc-QF-driven Tom expression. B. Loss of GAL80 results in de-repression of act-GAL4-driven GFP expression. C-C’. An example of a wing disc with a Tom clone. This disc also expresses en-GAL4>GFP that was used to express RNAi and transgenes in the posterior compartment of the disc. The genotype was en-GAL4>UAS-GFP, psc-QF>Tom, QS^5A^/+ from the cross of en-GAL4>UAS-GFP, psc-QF>Tom, QS^5A^/CyO-GFP with w^1118^. Larvae were treated as in panel E but without the temperature shift. D. An example of a wing disc with GFP clones. The genotype was GAL80^5B^/+; Act-GAL4, UAS-GFP/+ from the cross of GAL80^5B^/GAL80^5B^ with Act-GAL4, UAS-GFP/CyO RFP Tb. Larvae were treated as in panel E but without the temperature shift. E. The experimental protocol used to express RNAi and transgenes conditionally in the focused screen. For experiments that do not involve transgene expression, the same protocol was followed but larvae were maintained at 25°C without a temperature shift. F. Chromosomal location of QS^5A^, QS^9B^, *mwh* and GAL80 studied here (not to scale). Scale bar = 120 microns in C-D.

Alternatively, homozygous mutants for *mwh*, which encodes an actin-bundling protein, may be less fit while the loss of QS, which has no known role in *Drosophila*, may be neutral and it is this difference in gene function rather than the amount of chromosome lost that explains the differences in IR-induced LOH incidence. To address these possibilities and to identify regulators of IR-induced LOH, we studied the LOH incidence of six additional insertion loci (Fig 1F) and performed a focused genetic screen.

## Results

### IR-induced LOH shows sex-dependence but locus-independence

We used 4000 Rads (R) of X-rays, a typical dose in *Drosophila* (for example, (Brodsky *et al.* 2004; Wells and Johnston 2012)) because it kills more than half of the cells in wing discs, yet larvae can still develop into viable adults. Thus, 4000R causes significant damage but is still compatible with recovery and regeneration. To ask if the distance from the chromosome tip matters, we studied QS^5A^ that is further from the telomere than QS^9B^ or *mwh* (Fig 1F). As for QS^9B^ (Brown *et al.* 2020), LOH for QS^5A^ is IR-dependent (Fig 2A-B). More important, QS^9B^ and QS^5A^ did not differ significantly in IR-induced LOH clone number (Fig 2A) or clone area (Fig 2B). In feeding 3^rd^ instar larval stage, cells of the wing disc that will ultimately form the adult structures are in a single layer columnar epithelium with uniform cell size. Therefore, clone area serves as a proxy for cell number as in previous studies (Brown *et al.* 2020). We normalize clone number and area to the total disc area so that we can control for variations in disc size/total cell number. We conclude that distance from the chromosome tip did not affect IR-induced LOH incidence.

**Fig 2.**
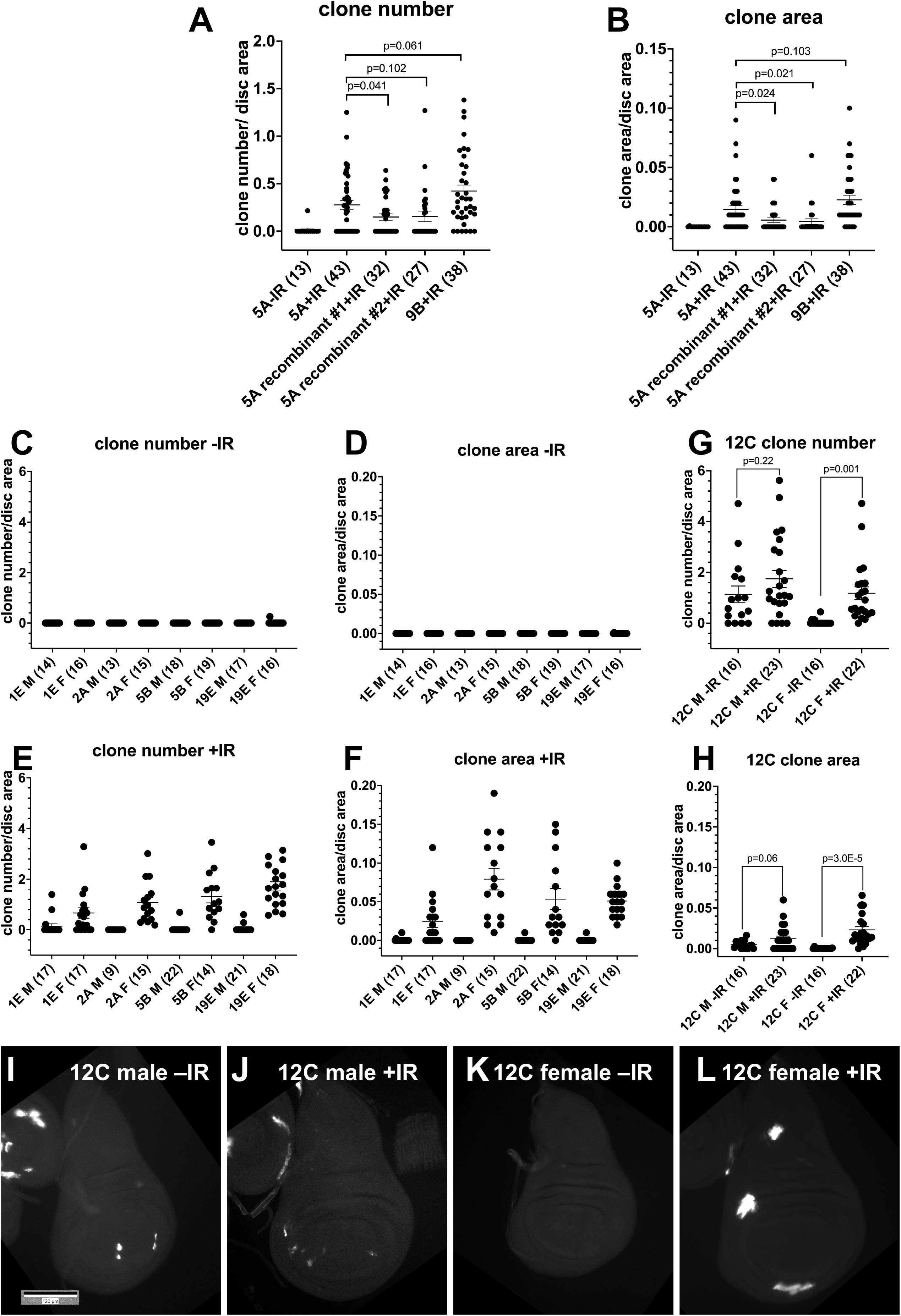
IR-induced LOH shows sex-dependence but locus-independence. Larvae were treated as in Fig 1E but without the temperature shift and wing discs were fixed and stained for DNA. The graphs show data are from at least two biological replicates per sample, with the total number of wing discs examined for each sample is shown in parentheses. Error bars represent mean±1 SEM. Scale bar = 120 microns. p-values were calculated using a 2-tailed t-test. (A-B) QS^9B^ and QS^5A^ are not significantly different in IR-induced LOH clone number (A) or clone area (B). (C-F) Quantification of GFP+ clone number (C-D) and clone area (E-F) with and without irradiation for four GAL80 insertions on the X chromosome. Male and female larvae were separated before dissection and processed separately. (G-H) Quantification of GFP+ clone number (G) and clone area (H) with and without irradiation for 12C6 GAL80 insertion on the X chromosome. Male and female larvae were separated before dissection and processed separately. (I-L) Representative examples of wing discs for the 12C insertion. The genotypes were: 9B = psc-QF>Tom/+; QS^9B^/+ that results from psc-QF>Tom; QS^9B^ X to w^1118^ 5A = psc-QF>Tom/QS^5A^ that results from psc-QF>Tom/CyO-GFP X QS^5A^/CyO-GFP (C-L) GAL80/+; Act-GAL4, UAS-GFP/+ that result from GAL80/GAL80 X Act-GAL4, UAS-GFP/CyO RFP Tb

The conclusion above is further supported by the study of GAL80 insertions at 5 loci along the X chromosome that was used before to detect spontaneous (without IR) LOH (Fig 1B,D,F) (Siudeja *et al.* 2015). For four of the five GAL80 insertions, wing discs from un-irradiated larvae show very few cells with GFP signal that is as bright as what is seen after IR (Fig 2C-D, Supplemental Fig 1). Occasional cells with faint GFP (arrow in Supplemental Fig 1A-A’’) are consistent with a recent report of spontaneous GAL80 ‘loss’ in larval brains through poorly understood mechanisms (Goupil *et al.* 2022). Importantly, we noticed in early experiments with GAL80 that approximately half of the irradiated discs lacked GFP expressing clones. This is different from what we saw with QS; for example, 32 of 38 QS^9B^ discs had Tom+ cells in Fig 2A ‘9B+IR’ dataset. QS insertions are on autosomes while GAL80 insertions are on the X chromosome. This led us to hypothesize a sex-dependence in GAL80 LOH, like what was reported for spontaneous LOH of X-linked loci (Siudeja *et al.* 2015). Analysis of wing discs from males and female larvae (see Methods) show that IR-induced increases in GFP+ cells were confined almost exclusively to females (Fig 2E-F).

The exception is the 12C insertion, which shows the greatest incidence of GFP+ cells without IR via unknown mechanisms, and a further increase after IR (Fig 2G-H). Even in this case, IR induced increase was significant only in the females (Fig 2I-L, quantified in G-H). Thus, IR-induced GAL80 LOH shows sex-dependence for all five insertions, similar to the case for spontaneous LOH (Siudeja *et al.* 2015). To our surprise, however, IR-induced LOH did not decrease with proximity to the centromere. This is unlike spontaneous LOH (Siudeja *et al.* 2015); GAL80^5B^ showed spontaneous LOH in 75% of adult *Drosophila* intestines while the corresponding number for GAL80^19E^ was 40%. In contrast, IR-induced LOH incidence was nearly identical for GAL80^5B^ and GAL80^19E^ (Fig 2E-F, p=0.193 for clone number and 0.861 for clone area). We conclude that IR-induced LOH shows sex-dependence (like spontaneous LOH) but chromosomal position independence (unlike spontaneous LOH). Possible explanations are in the Discussion.

### A focused screen identified E2F1 as a regulator of LOH

Results so far address cis-acting determinants, namely, chromosome position of the LOH loci and the presence of a homologous X chromosome. To identify trans-acting regulators, we performed a focused screen through known regulators of DNA damage and stress responses, cell growth, cell survival and cell competition (Supplemental Table 1), using published constructs that have been shown to produce a phenotype, when possible. We used QS in this screen so that we can use the GAL4/UAS system in parallel to knock down/overexpress genes. We used QS^9B^ because it shows more robust IR-induced LOH incidence than QS^5A^.

Transgenes were expressed in the posterior (P) half of wing discs using the protocol in Fig 1E. As stated above, we saw no Tom clones without IR; therefore, only +IR data are shown (Fig 3 and Supplemental Fig 2). Three controls were used: the TRiP III line that was the recipient of many RNAi constructs and RNAi for the w gene (both as controls for RNAi) and w^1118^ (control for overexpression or OE). After normalizing to compartment area to control for differences in compartment size, IR-induced LOH clone number or area are not significantly different between A and P compartments in all three controls (Fig 3A-B and Supplemental Fig 2). Most genes tested did not affect IR-induced LOH including those with known roles in cell competition such as duox and Xrp1 (Supplemental Fig 2). This group is of interest because a prior publication suggested that cells with segmental aneuploidy, a possible mechanism for LOH generation, may be eliminated by cell competition (Mcnamee and Brodsky 2009). We interpret negative results with caution, however, because our protocol for conditional and transient expression (Fig 1E) may not deplete gene function sufficiently even if the same construct produced a phenotype under different expression conditions before.

**Fig 3.**
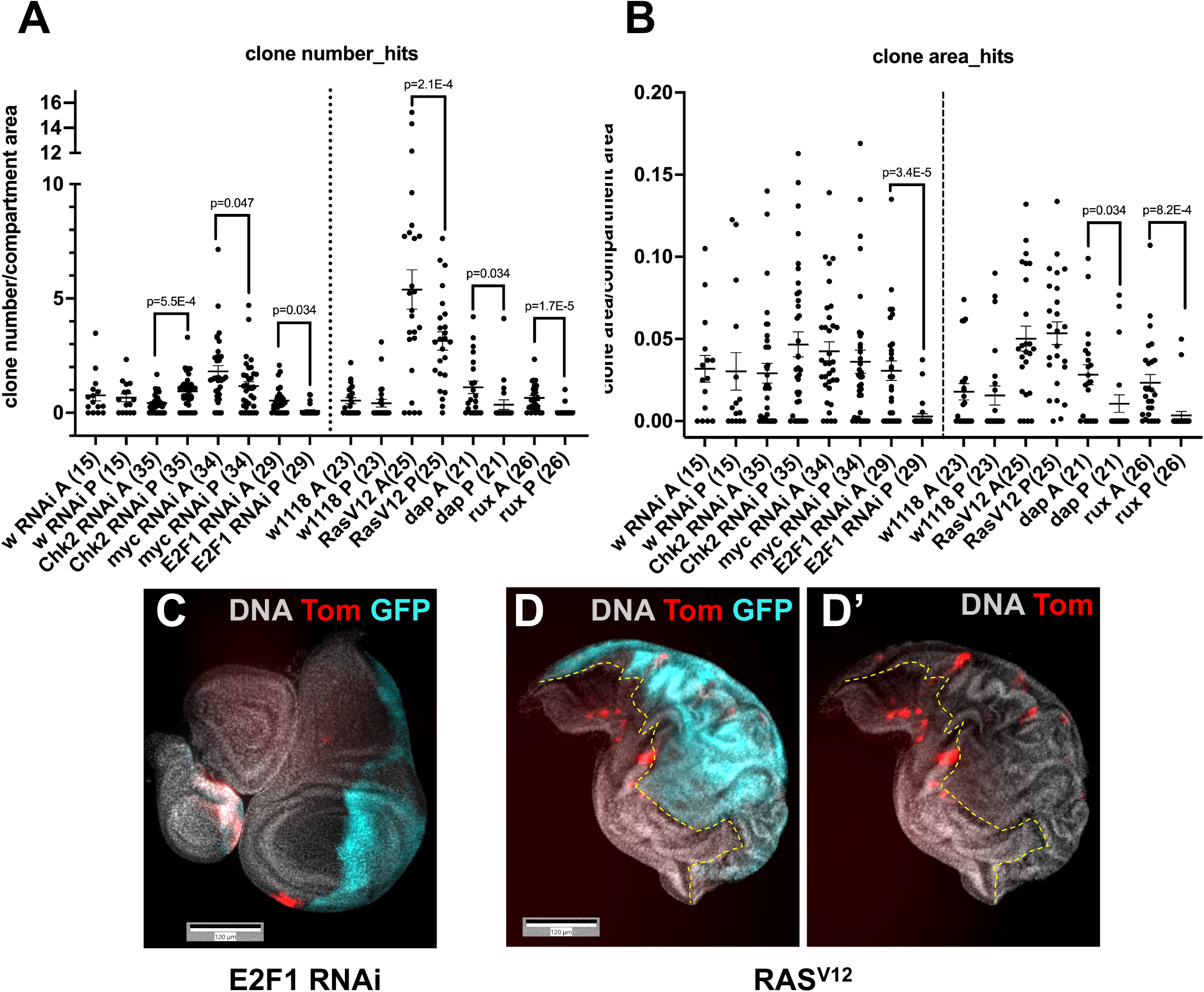
A focused screen identified E2F1 as a regulator of LOH. Wing discs from 3^rd^ instar larvae were dissected 3 days after irradiation, fixed and stained for DNA. Larvae were treated as in Fig 1E. The graphs show data from at least two biological replicates per sample, with the total number of wing discs examined per sample shown in parentheses. The mean±1 SEM is shown for each sample. p-values were calculated using a paired 2-tailed t-test. (A-B) Results of the focused screen showing clone number (A) and clone area (B). RNAi against w is shown as the control for RNAi constructs and w^1118^ is shown as the control for overexpression constructs. (C-D’) Representative examples of wing discs expressing E2F1^RNAi^ or RAS^V12^ in the P compartment (GFP+).

The screen found mild effects with RNAi against Chk2 or Myc and with overexpression of oncogenic Ras^V12^ (Fig 3A-B). Specifically, Chk2 RNAi increased LOH clone number while Myc RNAi and Ras^V12^ decreased LOH clone number. The effect sizes, however, were small (1.5 to 2-fold) and clone area was not significantly affected, suggesting compensating changes in the size of individual clones. In addition, expression of Ras^V12^ in the P compartment produced epithelial folds in both compartments in irradiated discs (Fig 3D-D’), making the quantification of clone area challenging. Because of this and small effect sizes, we have not followed up on Chk2, Myc and Ras.

In contrast, RNAi against E2F1 profoundly affected both LOH clone number and clone area with a reduction of the former by 6-fold and the latter by more than 10-fold (Fig 3C, quantified in A-B). *Drosophila* E2F1, like its mammalian homologs, promotes cell proliferation by promoting G1/S transition via activation of cyclin E expression and G2/M transition via activation of Cdc25^String^ (Reis and Edgar 2004). E2F1 also promotes IR-induced apoptosis in the wing discs when p53 has been depleted; that is, its role in apoptosis is negligible in a wild type background (Wichmann *et al.* 2006). Therefore, we addressed the more likely possibility that the requirement for E2F1 in LOH clone formation/maintenance is through its role in the cell cycle. To this end, we expressed Cdk inhibitors Dap and Rux to inhibit G1/S and G2/M transitions, respectively, using the same temperature shift protocol as in Fig 1E. Both Dap and Rux phenocopied E2F1 RNAi (Fig 3A-B), supporting the idea that the role of E2F1 is to promote the proliferation of cells with LOH.

We normalize LOH clone number and area to the compartment area. If E2F1 depletion or Dap/Rux overexpression in the P compartment affected all cells within this compartment equally regardless of their LOH status, we should see little change in <u>normalized</u> LOH clone number or area. Yet, we see multiple-fold reductions by both measures. We infer that manipulations of cell cycle regulators have a more severe effect on cells with LOH than on cells in the rest of the P compartment. Greater sensitivity of LOH cells to cell cycle inhibition is a new finding.

The manipulations thus far alter gene expression in the whole compartment and identified differential effects on LOH versus non-LOH cells. To address cell-autonomous needs in LOH cells more directly, we took advantage of QS loss and consequent QF de-repression to express QUAS-transgenes and assayed their effect. This protocol has the added advantage of identifying needs AFTER the clones have formed and have initiated QF-mediated gene expression. As proof-of-concept, we used a published QUAS-rpr transgene (Perez-Garijo *et al.* 2013) to express this pro-apoptotic gene specifically in LOH clones, with the expectation that such clones will be lost to cell death. The results show the system works as expected; QUAS-rpr reduced both clone number and clone area to near completion (Fig 4A-B, quantified in E-F).

**Fig 4.**
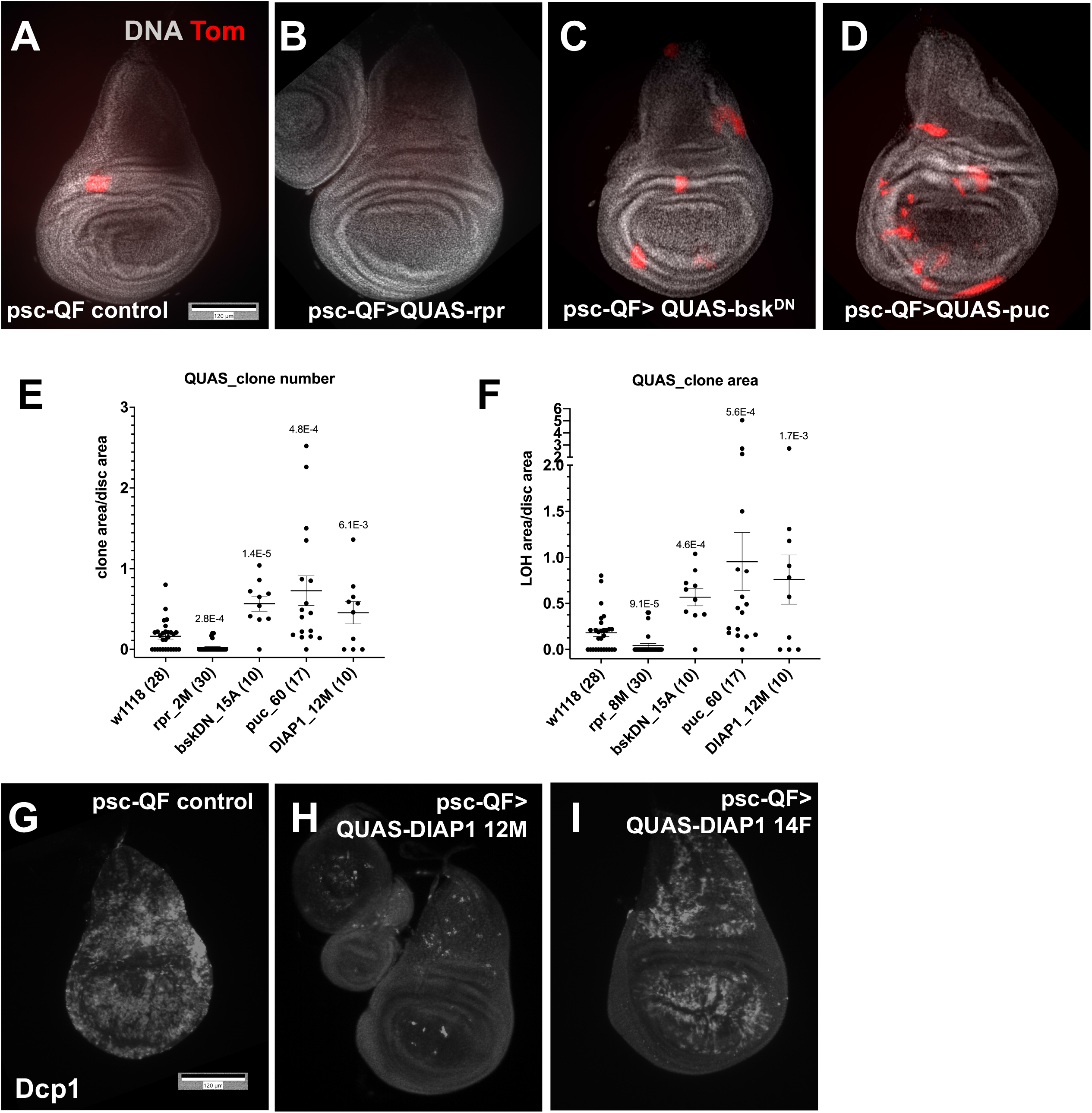
Cell autonomous requirements for LOH. (A-D) Larvae were treated as in Fig 1E but without the temperature shift and wing discs were fixed and stained to visualize DNA. The discs shown are from irradiated larvae. (E-F) Quantification of Tom+ clone number (E) and clone area (F) in irradiated discs. The graphs show data from at least two biological replicates per sample, with the total number of wing discs examined per sample shown in parentheses. The mean±1 SEM is shown for each sample. p-values relative to w^1118^ controls were calculated using a 2-tailed t-test. (G-I) Larvae were cultured and irradiated as in (A-F) but wing discs were dissected 4h after IR, fixed and stained for DNA and with an antibody against cleaved active Dcp1 as a surrogate marker for apoptosis. Scale bars = 120 microns. The genotypes are as shown below. Stock information is in Supplemental Table 1. (A,G) Control = psc-QF>Tom/+; QS^9B^/+ that results from psc-QF>Tom; QS^9B^ X to w^1118^ (H-I) psc-QF>Tom/QUAS-transgene; QS^9B^/+ (B-D) psc-QF>Tom/+; QS^9B^/QUAS-transgene

Using QUAS-transgenes, we tested the role of JNK and apoptotic signaling in maintaining cells with LOH. We first assessed transgene functionality by ubiquitous expression with psc-QF (without QS) and assaying for expected phenotypes. For rpr, dominant negative bsk (JNK) and puc (phosphatase and JNK inhibitor), we found lines that produced significant lethality (Supplemental Table 2). For *Drosophila* Inhibitor of Apoptosis Protein 1 (DIAP1), even the strongest effect on lethality was modest, so we assayed for inhibition of apoptosis instead. Of three lines tested, 12M reduced IR-induced cleaved Dcp1 signal to near completion (Fig 4H), 14F showed a modest effect (Fig 4I), and 3M showed variable effect from disc to disc. Therefore, QUAS-DIAP1 12M line was used in subsequent experiments. The results show that cell-autonomous inhibition of JNK or apoptosis in cells with LOH led to a significant increase in both Tom clone number and area. We conclude that in cells that survived LOH long enough to activate QUAS-transgenes, stress signaling continues to operate to eliminate them.

## Discussion

We report here studies on the role of chromosomal position and trans-acting regulators on IR-induced LOH in *Drosophila* larval wing discs. We found that position along the chromosome makes little difference while the presence of a homologous chromosome is required. We identified E2F1 as a positive regulator of LOH clone incidence and propose that cells with LOH are more sensitive to inhibition of cell proliferation than their neighbors. And we identified JNK signaling and apoptosis as cell-autonomous mechanisms that eliminate cells that survived LOH induction. This contrasts with the previous finding that JNK plays only a backup role in reducing IR-induced *Minute* (M/+) bristles, a role that becomes apparent only in p53 mutant background (McNamee and Brodsky 2009).

Spontaneous LOH in the adult *Drosophila* intestine is attributed to mitotic recombination, based on several lines of evidence. First, sex-dependence is interpreted as arising from recombination-based mechanisms that require the presence of a homologous chromosome such as crossing over, gene conversion or break-Induced replication (Siudeja *et al.* 2015). Second, adult guts with spontaneous GAL80^5B^ LOH was about twice as frequent as those with spontaneous GAL80^19E^ LOH. An increase in LOH with distance from the centromere is predicted for recombination-based mechanisms because a distal locus would experience more frequent exchanges between homologous chromosomes. This study noted, however, that the frequency of spontaneous LOH at 19E was higher than predicted from meiotic maps and higher than observed for a centromere-proximal locus on chromosome 3R, suggesting additional factors at play. Third, whole genome sequencing confirmed that spontaneous LOH of chromosome 2 loci in the adult intestine occur through mitotic recombination (zouabi *et al.* 2022).

Our finding that IR-induced LOH clone number and area were nearly identical for GAL80^5B^ and GAL80^19E^ cannot be explained by recombination-based mechanisms. Classical studies by Baker and others monitored several adult visible markers such as *yellow (y)* and *forked (f)* in heterozygous adult females that were irradiated as larvae or pupae with 1000 R from a Co-60 source (Baker *et al.* 1978). In adult female abdomens monitored for *y* or *f*, both X-linked genes, 40% of LOH clones induced by IR (i.e., the difference between -/+IR samples) consisted of 40% twin clones (likely products of mitotic recombination) while the remaining 60% were single clones. Examination of single clones that arose spontaneously (without radiation) in the same study found that 20% of cells in such clones showed *Minute* bristles, which result from haploinsufficiency of loci encoding ribosomal proteins. The authors suggest that single LOH clones arose by aneuploidy that turned wild type (+/+) cells into heterozygotes for ribosomal protein genes (M/+). In another set of studies, McNamee and Brodsky used the *Minute* bristle phenotype and assayed gain-of-heterozygosity, from +/+ to M/+, rather than loss-of-heterozygosity, in bristles from adults that result from irradiated larvae (McNamee and Brodsky 2009). Recombination-based mechanisms induced by IR would not be detectable in this assay as it would change +/+ to +/+. Regardless, taken together, these results suggest that both aneuploidy and recombination-based mechanisms operate to produce IR-induced LOH in tissues that produce the adult body wall. Our data suggest a greater contribution by non-recombination-based mechanisms, namely aneuploidy.

Studies by Baker, Brodsky and colleagues did not distinguish among various types of aneuploidies, which could include loss of whole or segments of chromosomes and terminal deletions. Over-expression of telomere capping protein HipHop has been shown to increase the survival of cells with terminal deletions that result from dividing experimentally induced dicentric chromosomes (Kurzhals *et al.* 2017). Overexpression of HipHop, however, did not increase the incidence of IR-induced LOH (Supplemental Fig 2), suggesting that terminal deletions may not be the mechanism. The finding that distance from the telomere did not affect the QS LOH incidence supports this idea. Instead, relevant types of aneuploidy may include X-ray-induced segmental losses that are well-documented in numerous studies in *Drosophila* and mammals (for example, (Pastink *et al.* 1987; Pastink *et al.* 1988; Eeken *et al.* 1994; Chick *et al.* 2005)). Such losses can explain our finding that LOH of X-linked loci requires a homologous chromosome. We suggest that without part or whole of the X chromosome in XY males, a cell may not survive to express GFP, reducing the level of LOH observed in males. In other words, presence of one X chromosome allows cells to survive with partial or complete loss of the other X chromosome.

## Materials and Methods

### Drosophila stocks and methods

*Drosophila* stocks used are in Supplemental Table 1 and include those with UAS-puc (Martin-Blanco *et al.* 1998), UAS-Wg (Lawrence *et al.* 1996) and recombinant chromosome 3 encoding tub-QS^9B^ and tub-GAL80^ts^ (Brown *et al.* 2020). QUAS lines are a gift from Ainhoa Pérez-Garijo and Hermann Steller (Perez-Garijo *et al.* 2013). Virgin females and males were crossed and cultured for three days on Nutri-Fly German Formula food (Genesee Scientific) at 25°C before egg collections. Eggs were collected and larvae were raised on Nutri-Fly Bloomington Formula food (Genesee Scientific) at 25°C unless otherwise noted. The cultures were monitored daily for signs of crowding, typically seen as ‘dimples’ in the food surface, and were split as needed. Larvae in food were placed in petri dishes and irradiated in a Faxitron Cabinet X-ray System Model RX-650 (Lincolnshire, IL) at 115 kVp and 5.33 rad/sec. Larval sex was determined using gonad size observed under a dissecting microscope; male gonads are large and free of the fat body while female gonads are hard to find (Kerkis 1931).

### Tissue preparation and imaging

Larval wing discs were dissected in PBS, fixed in 4% para-formaldehyde (PFA) in PBS for 30 min, and washed three times with PBTx (0.1% Triton X-100). The discs were stained with 10 μg/ml Hoechst33342 in PBTx for 5 min, washed 3 times, and mounted on glass slides in Fluoromount G (SouthernBiotech). Wing discs were imaged on a Leica DMR compound microscope using a Q-Imaging R6 CCD camera and Ocular or Micro-Manager software.

### Statistical Analysis

For sample size justifications, we used a simplified resource equation from (Charan and Kantharia 2013); E = total number of animals − total number of groups, where E value of 10-20 is considered adequate. When we compare two groups (-IR and +IR, for example, where the former is the control group and the latter is the treated group), n = 6 per group, or E = 11, would be adequate. All samples subjected to statistical analysis meet or exceed this criterion. To compute p-values, 2-tailed Student t-tests were used.

## Data and Reagent Statement

Drosophila stocks are available upon request. The authors affirm that all data necessary for confirming the conclusions of the article are present within the article, figures, and tables.

## Acknowledgments

We thank the Perrimon, Bardin and Steller labs for fly stocks. Additional stocks from the Bloomington *Drosophila* Stock Center (NIH P40OD018537) and Vienna *Drosophila* Resource Center were used in this study.

## Figure and Table Legends

**Supplemental Figure 1.**
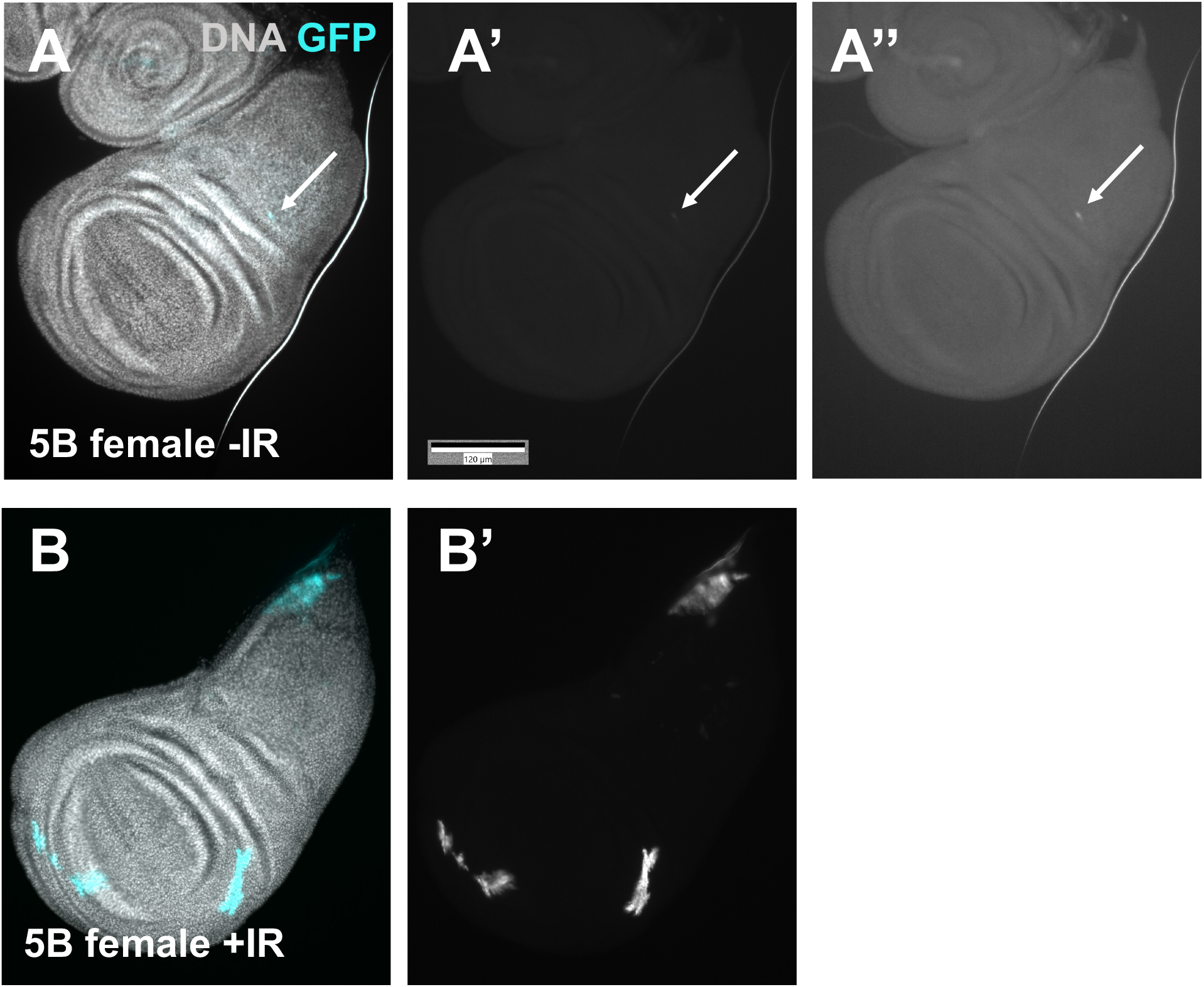
Spontaneous GFP is less bright than IR-induced GFP. Larvae were treated as in Fig 1E but without the temperature shift and wing discs were fixed and stained to visualize DNA. The genotype of the larvae was GAL805B/+; Act-GAL4, UAS-GFP/+ from the cross of GAL805B/GAL805B with Act-GAL4, UAS-GFP/CyO RFP Tb. (A-A’’’) A disc from an unirradiated larva showing one faint GFP spot (arrow). (B-B’) A disc from an irradiated larva showing bright GFP spots. A’ and B’ are shown with similar brightness and contrast range in ImageJ so that signals would be directly comparable. A’’ is shown with a narrower range to allow the viewer to see the faint GFP spot.

**Supplemental Figure 2.**
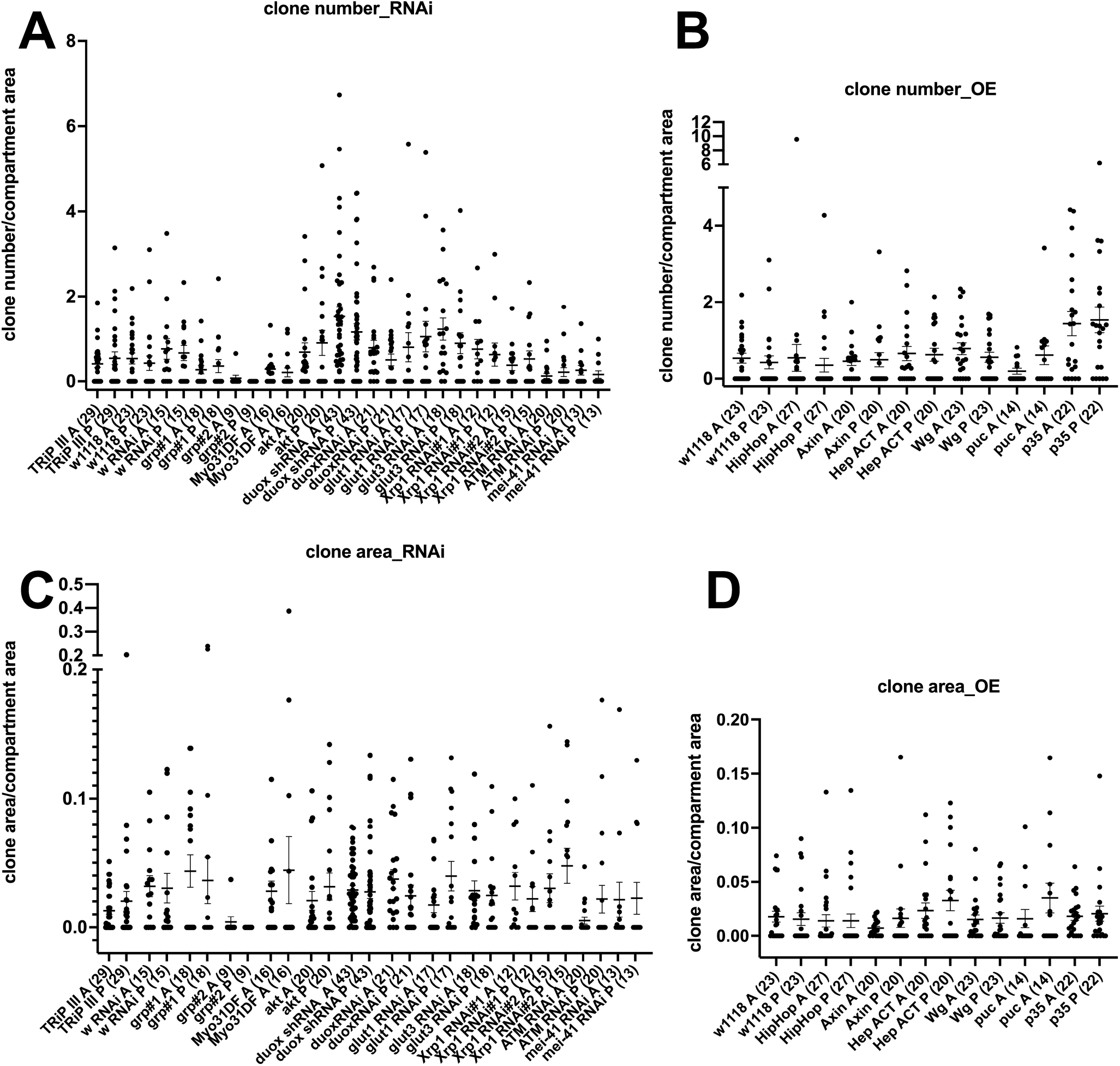
A focused screen for modulators of IR-induced QS9B LOH. Wing discs from 3rd instar larvae were dissected 3 days after irradiation, fixed, and stained for DNA. Larvae were treated as in Fig 1E. The graphs show data from at least two biological replicates per sample, with the total number of wing discs examined per sample shown in parentheses. The mean±1 SEM is shown for each sample. Because A and P compartments are of different sizes, clone numbers and clone areas have been normalized to compartment area. p-values between A and P samples for each genotype (none <0.05) were calculated using a paired 2-tailed t-test. TRiP III (background stock for most RNAi lines used) and RNAi against w served as controls for RNAi constructs and w1118 served as the control for overexpression constructs. (A-B) LOH clone numbers for RNAi (A) and overexpression (OE,B) constructs. (C-D) LOH clone areas for RNAi (C) and overexpression (OE,D) constructs.

**Supplemental Table 1.**
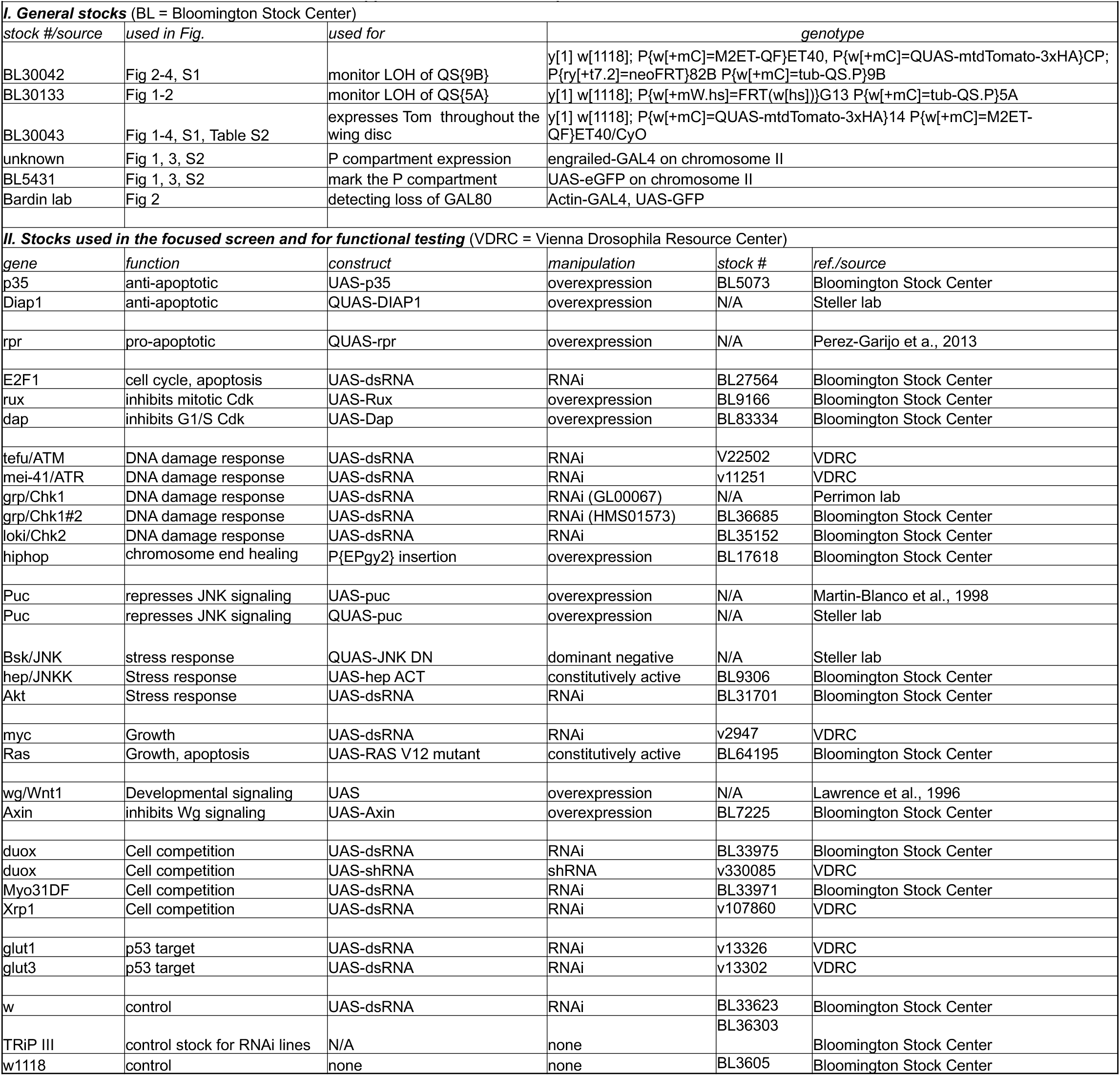
Drosophila stocks used in this work.

**Supplemental Table 2.**
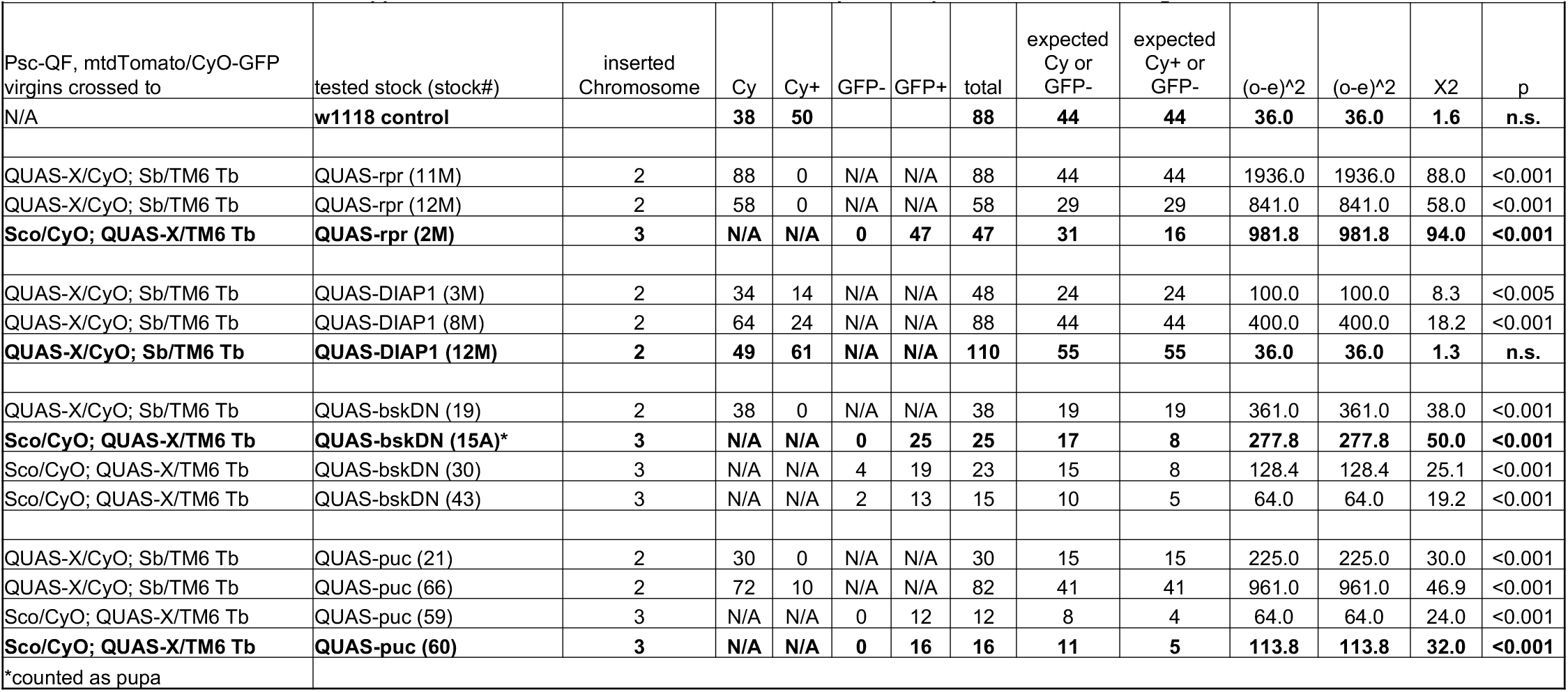
Χ^2^ test for survival after ubiquitous expression of QUAS transgenes. For chromosome 2 lines, CyO balancer was floating so Cy+ males were used. Male QUAS flies were crossed to virgins of the genotype psc-QF>UAS-Tomato/CyO-GFP. Embryos were collected overnight. CyO and CyO+ progenies were counted until no new adults eclosed for a 24 h period. Χ^2^ test was used to assess deviations from the 1:1 expected Mendelian ratio. For insertions on chromosome 3, Tb+ pupae were sorted and scored as GFP+ or GFP-before eclosion was monitored. Χ^2^ test was used to assess deviations from the 2GFP- (Sco/QF>Tom and QF>Tom/CyO):1GFP+ (Sco/CyO-GFP) expected Mendelian ratio.

| Supplemental Table 2. X2 test for survival after ubiquitous expression of QUAS transgenes. |  |  |  |  |  |  |  |  |  |  |  |  |  |
| --- | --- | --- | --- | --- | --- | --- | --- | --- | --- | --- | --- | --- | --- |
| Psc-QF, mtdTomato/CyO-GFP virgins crossed to | tested stock (stock#) | inserted Chromosome | Cy | Cy+ | GFP- | GFP+ | total | expected Cy or GFP- | expected Cy+ or GFP- | (o-e)^2 | (o-e)^2 | X2 | p |
| N/A | w1118 control |  | 38 | 50 |  |  | 88 | 44 | 44 | 36.0 | 36.0 | 1.6 | n.s. |
| QUAS-X/CyO; Sb/TM6 Tb | QUAS-rpr (11M) | 2 | 88 | 0 | N/A | N/A | 88 | 44 | 44 | 1936.0 | 1936.0 | 88.0 | <0.001 |
| QUAS-X/CyO; Sb/TM6 Tb | QUAS-rpr (12M) | 2 | 58 | 0 | N/A | N/A | 58 | 29 | 29 | 841.0 | 841.0 | 58.0 | <0.001 |
| Sco/CyO; QUAS-X/TM6 Tb | QUAS-rpr (2M) | 3 | N/A | N/A | 0 | 47 | 47 | 31 | 16 | 981.8 | 981.8 | 94.0 | <0.001 |
| QUAS-X/CyO; Sb/TM6 Tb | QUAS-DIAP1 (3M) | 2 | 34 | 14 | N/A | N/A | 48 | 24 | 24 | 100.0 | 100.0 | 8.3 | <0.005 |
| QUAS-X/CyO; Sb/TM6 Tb | QUAS-DIAP1 (8M) | 2 | 64 | 24 | N/A | N/A | 88 | 44 | 44 | 400.0 | 400.0 | 18.2 | <0.001 |
| QUAS-X/CyO; Sb/TM6 Tb | QUAS-DIAP1 (12M) | 2 | 49 | 61 | N/A | N/A | 110 | 55 | 55 | 36.0 | 36.0 | 1.3 | n.s. |
| QUAS-X/CyO; Sb/TM6 Tb | QUAS-bskDN (19) | 2 | 38 | 0 | N/A | N/A | 38 | 19 | 19 | 361.0 | 361.0 | 38.0 | <0.001 |
| Sco/CyO; QUAS-X/TM6 Tb | QUAS-bskDN (15A)* | 3 | N/A | N/A | 0 | 25 | 25 | 17 | 8 | 277.8 | 277.8 | 50.0 | <0.001 |
| Sco/CyO; QUAS-X/TM6 Tb | QUAS-bskDN (30) | 3 | N/A | N/A | 4 | 19 | 23 | 15 | 8 | 128.4 | 128.4 | 25.1 | <0.001 |
| Sco/CyO; QUAS-X/TM6 Tb | QUAS-bskDN (43) | 3 | N/A | N/A | 2 | 13 | 15 | 10 | 5 | 64.0 | 64.0 | 19.2 | <0.001 |
| QUAS-X/CyO; Sb/TM6 Tb | QUAS-puc (21) | 2 | 30 | 0 | N/A | N/A | 30 | 15 | 15 | 225.0 | 225.0 | 30.0 | <0.001 |
| QUAS-X/CyO; Sb/TM6 Tb | QUAS-puc (66) | 2 | 72 | 10 | N/A | N/A | 82 | 41 | 41 | 961.0 | 961.0 | 46.9 | <0.001 |
| Sco/CyO; QUAS-X/TM6 Tb | QUAS-puc (59) | 3 | N/A | N/A | 0 | 12 | 12 | 8 | 4 | 64.0 | 64.0 | 24.0 | <0.001 |
| Sco/CyO; QUAS-X/TM6 Tb | QUAS-puc (60) | 3 | N/A | N/A | 0 | 16 | 16 | 11 | 5 | 113.8 | 113.8 | 32.0 | <0.001 |
| *counted as pupa |  |  |  |  |  |  |  |  |  |  |  |  |  |

